# Culture of salivary methanogens assisted by chemically produced hydrogen

**DOI:** 10.1101/734210

**Authors:** CO Guindo, E Terrer, E Chabrière, G Aboudharam, M Drancourt, G Grine

## Abstract

Methanogen cultures require hydrogen produced by fermentative bacteria such as *Bacteroides thetaiotaomicron* (biological method). We developed an alternative method for hydrogen production using iron filings and acetic acid with the aim of cultivating methanogens more efficiently and more quickly (chemical method). We developed this new method with a reference strain of *Methanobrevibacter oralis*, compared the method to the biological reference method with a reference strain of *Methanobrevibacter smithii* and finally applied the method to 50 saliva samples. Methanogen colonies counted using ImageJ software were identified using epifluorescence optical microscopy, real-time PCR and PCR sequencing. For cultures containing the pure strains of *M. oralis* and *M. smithii*, colonies appeared three days postinoculation with the chemical method versus nine days with the biological method. The average number of *M. smithii* colonies was significantly higher with the chemical method than with the biological method. There was no difference in the delay of observation of the first colonies in the saliva samples between the two methods. However, the average number of colonies was significantly higher when using the biological method than when using the chemical method at six days and nine days postinoculation (Student’s test, p = 0.005 and p = 0.04, respectively). The chemical method made it possible to isolate four strains of *M. oralis* and three strains of *M. smithii* from the 50 saliva samples. Establishing the chemical method will ease the routine isolation and culture of methanogens.

## INTRODUCTION

Methanogenic archaea (referred to herein as methanogens) are acknowledged members of the digestive tract microbiota, and they have been detected by PCR-based methods and cultured from the oral cavity and the stools of apparently healthy individuals (1). More particularly, *Methanobrevibacter oralis, Methanobrevibacter smithii* and *Methanobrevibacter massiliense* have been isolated from the oral cavity, whereas *M. smithii, Methanosphaera stadtmanae, Methanomassiliicoccus luminiyensis, Methanobrevibacter arboriphilicus, M. oralis, Ca*. Methanomethylophilus alvus and *Ca*. Methanomassiliicoccus intestinalis have been isolated from stools (1). Accordingly, we previously showed that virtually all apparently healthy individuals would carry methanogens in the digestive tract microbiota (2). This observation indeed corroborated the pivotal role of methanogens, which detoxify hydrogen produced by bacterial fermentations into methane (3, 4).

Moreover, methanogens are increasingly implicated in diseases; their presence or absence is associated with dysbioses such as those observed in the gut microbiota in cases of chronic constipation (5), obesity (6) and colonic diseases including ulcerative colitis, Crohn’s disease and colorectal cancer (7–9), in the vaginal microbiota in cases of vaginosis (10, 11), or their presence in anaerobic pus abscesses such as brain abscesses (12, 13) and muscular abscesses (14).

As is usual in clinical microbiology, isolation and culture of methanogens is the gold standard to assess the detection of living methanogens in microbiota and in pathological clinical specimens collected by puncture or biopsy (1). The routine application of methanogen culturing is hampered by the fact that methanogens are strictly aero-intolerant microbes and require hydrogen for culturing (15, 16). To facilitate the isolation and culture of methanogens from a routine perspective, we previously designed a new process in which methanogens are cultivated in the presence of hydrogen-producing *Bacteroides thetaiotaomicron* (17, 18). Here, we tested the conditions to replace the biological production of hydrogen with a chemical production of hydrogen and applied it to isolate methanogens from saliva as a proof-of-concept.

## MATERIALS AND METHODS

### Chemical production of hydrogen

The immediate chemical production of hydrogen resulting from the oxidation of iron by a weak acid has long been known (19). We used acetic acid in our experiment because of its ability to oxidize iron, resulting in sufficient production of hydrogen (20). We used two solutions for our experiments, namely, SAB culture broth (21) and distilled water (Fresenius, Bad Homburg voor Hoehe, Germany) with pH values of 7.3 and pH values of 7, respectively. We used 5 mL of each solution in our experiments. The experiments were performed in 10 Hungate tubes (Dominique Dutscher, Brumath, France) for each solution. Iron filings (Amazon, Clichy, France) were used at increasing quantities: 0. 5 g for tube n°1, 1 g for tube n°2, and a 0.25 g increase from tube n°3 to tube n°10. The amount of acetic acid (VWR International, Pennsylvania, USA) used was 100 μL for tube n°1 to tube n°7 and 50 μL for tube n°8 to tube n°10. We first put the iron filings in the tubes, followed by the solution (SAB broth or water) and then the acetic acid. We used three negative control Hungate tubes, one tube with iron filings in SAB culture medium, another tube with iron filings in distilled water and finally a third tube containing iron filings alone. Gas chromatography was then performed using a Clarus 580 FID chromatograph (PerkinElmer, Villebon-sur-Yvette, France) to measure hydrogen production in the different tubes, and the pH was monitored (Fisher Scientific, Illkirch, France).

### Culture of methanogen strains using the chemical method for hydrogen production

*M. oralis* CSUR P9633, a human saliva isolate (22), was used for the development of the chemical method. We used the mini-double-chamber flask technique, derived from the one previously described (21). SAB agar plates (5-cm plates, Fisher Scientific) inoculated with 200 μL of a *M. oralis* suspension at 10^7^ colony-forming units (CFUs) was placed in the upper compartment of the mini-double-chamber flask, which was sealed with parafilm to ensure an anaerobic atmosphere. We then put 1 g of iron filings and 100 μL of acetic acid in 10 mL of distilled water in the lower compartment of the mini-double-chamber flask in a microbiological safety station. We used an SAB agar plate inoculated with 200 μL of sterile phosphate buffered saline (PBS) (Fisher Scientific) as a negative control. We then incubated the plates at 37°C for nine days with visual inspection on day 3, day 6 and day 9 postinoculation.

Then, the growth of *M. smithii* CSUR P9632, a human stool isolate, was compared using the chemical method with the reference biological method using a double-chamber system as previously described for the aerobic culture of methanogens (17). For the chemical method, we put 1.5 g of iron filings and 150 μL of acetic acid in 200 mL of distilled water in the lower compartment instead of *Bacteroides thetaiotaomicron*, which was used in the biological method. SAB agar plates inoculated with 200 μL of a *M. smithii* suspension at 10^6^ CFU or 200 μL of PBS for the negative controls were placed in the upper compartment.

The pH of the SAB agar plates was measured at day 3, day 6 and day 9 postinoculation using pH indicator strips (Macherey-Nagel SARL, Hœrdt, France). We placed the pH indicator strip directly on the SAB agar plate for two minutes. We then took the value corresponding to the color obtained on the strip from the values indicated by the supplier. The pH and the redox potential of the broth in the lower chamber were measured at day 3, day 6 and day 9 postinoculation using the Accumet™ AE150 apparatus (Fisher Scientific). The probe was rinsed thoroughly with distilled water and then immersed in the lower chamber until the device displayed the value on the reading screen. The rinsing step was performed after each use to avoid any error in reading the device. We used the pH and redox potential of distilled water alone and the SAB culture broth alone as controls. We also measured the redox potential of the upper compartment. For the two methods, we used one double-chamber system, in which we put one Petri plate with 10 mL of distilled water and one Petri plate with 10 mL of the SAB culture broth. These double-chamber systems were then incubated at 37°C, and the redox potential was measured in each of the Petri dishes at day 3, day 6 and day 9 postinoculation. The effectiveness of the two methods for the growth of methanogens was compared by observing the appearance of the first colonies as well as the average number of colonies at day 3, day 6 and day 9 postinoculation. Colonies were confirmed by autofluorescence optical microscopy as follows. Briefly, the colony deposited on a microscopy slide was observed at 63X using an epifluorescence microscope (Leica DMI 3000, Wetzlar, Germany) and at 100X magnification using another epifluorescence microscope (Leica DMI 6000). Colonies were enumerated at day 3, day 6 and day 9 postinoculation by focusing the image of the entire 5-cm plate on a black background using a portable camera without flash (Lenovo L38011, Beijing, China). Numbered images were analyzed and counted using ImageJ version 8 (https://imagej.nih.gov/ij/download.html) (Wayne Rasband, Java, National Institutes of Health, USA) as follows: each image was imported into ImageJ software and analyzed in a first step using the blue, diamidino phenylindole filter to normalized the image. In a second step, the normalized image was analyzed using the green fluorescein isothiocyanate filter and saved in the Joint Photographic Experts Group format. We standardized the images using the elliptical selection tabulation of the software. We then proceeded to manually count the enlarged colonies using the magnifying glass and the multipoint tabulation of the software.

### Comparison of the two methods on saliva samples

The study was previously approved by the Ethics Committee of the IHU Méditerranée infection under n°2016-011. After having informed and collected the consent of the participants, we collected saliva samples from 50 people, including 25 tobacco smokers (15 men, 10 women, median age of 30) and 25 nonsmokers (12 men, 13 women, median age of 28). The collection and processing of samples were performed as previously described (22). We used SAB agar plates inoculated with 200 μL of PBS as negative controls and then proceeded in the same manner as described above for the *M. smithii* strain.

### Molecular analysis

PCR was performed to confirm the identity of the colonies. Colonies were suspended in 200 µL of ultrapure water (Fisher Scientific), and a sonication step was performed for 30 minutes. DNA was then extracted with the EZ1 Advanced XL Extraction Kit (QIAGEN, Hilden, Germany) using 200 µL as the sample volume and 200 µL as the elution volume. Gene amplification and PCR sequencing were performed as previously described (22–24). Real-time PCR analyses were performed as previously described (25).

## RESULTS AND DISCUSSION

We report on the use of chemically produced hydrogen for the isolation and culture of methanogens as an alternative to the currently used biological method (17). The method we report is based on the production of hydrogen resulting from the chemical reaction between iron filings and a weak acid such as acetic acid. All reported results were validated by negative controls that remained negative and identification of colonies by autofluorescence and PCR-based analyses.

We first determined an optimal balance between the concentration of iron filings and acetic acid to maintain the production of hydrogen over several days at a relative concentration compatible with the isolation and culture of methanogens. These preliminary experiments enabled the development of a safe and controlled production of hydrogen that was shown to be efficient in the culture of *M. oralis* and *M. smithii* reference strains.

In the first step, we measured the average redox potential and pH values of the various media investigated here. The pH and redox potential of the controls consisting of distilled water and SAB broth culture medium were 7 and + 81.3 mV and 7.3 and −11.9 mV, respectively. After incubation at 37°C for 3, 6 and 9 days, these values were 6.98, 6.92 and 6.87 units and +64.6 mV, +67.3 mV and +79.9 mV for distilled water; and 6.83, 6.78 and 6.27 units and +11.1 mV, +12.9 mV and +14.8 mV for the SAB broth culture medium, respectively (Table 1). These results agree with previous observations that most types of water, including tap water and bottled water, are oxidizing agents, and the value of their redox potential is positive, ≥ 20 mV (26). The addition of antioxidants in the SAB medium makes this culture medium reductive. Then, the pH and redox potential of distilled water enriched with iron filings and acetic acid (chemical method) were measured at 5.23, 5.16 and 5.08 units and +109.77 mV, +108.6 mV and +112.7 mV, respectively (Table 2). These values were 5.8, 5.25, and 5.24 units and +113.8 mV, +113.8 mV and +114.5 mV at day 3, day 6 and day 9 for the SAB culture medium incubated with *B*. *thetaiotaomicron*, respectively (Table 2). These data indicated that the cultivation of *B. thetaiotaomicron* in the SAB medium releases fermenting compounds such as CO2, hydrogen and acetate, which could oxidize the SAB medium (26). Then, we measured these values in the Petri dishes placed in the upper compartment of the double-chamber system, which hosted the methanogen culture (Fig. 1). The redox potential values were −28.1 mV, −81.2 mV and −108.9 mV at day 3, day 6 and day 9 for distilled water, respectively, and +13.5 mV, −116.5 mV and −77.5 mV at day 3, day 6 and day 9 for the SAB culture broth in the chemical method, respectively; and for the biological method, the redox potential values were −4.5 mV, −66 mV and −91.6 mV at day 3, day 6 and day 9 for distilled water, respectively, and +19.1 mV, −7.1 mV and −34.9 mV at day 3, day 6 and day 9 for the SAB culture broth in the biological method, respectively (Table 3). These data indicated that the hydrogen, being a reducer released into the bottom compartment, reaches the upper compartment and reduces the SAB medium or the water placed into Petri dishes in the upper compartment (27).

**Table 1.** Controls of pH and redox potential.

| Controls | Redox potential (mV) |  |  |  | pH |  |  |  |
| --- | --- | --- | --- | --- | --- | --- | --- | --- |
|  | D0 | D3 | D6 | D9 | D0 | D3 | D6 | D9 |
| Distilled water | +81.3 | +64.6 | +67.3 | +79.9 | 7 | 6.98 | 6.92 | 6.87 |
| SAB medium | -8.7 | +11.1 | +12.9 | +14.8 | 7.3 | 6.83 | 6.78 | 6.27 |

**Table 2.** pH and redox potential of the lower compartment.

| Methods | Redox potential (mV) |  |  | pH |  |  |
| --- | --- | --- | --- | --- | --- | --- |
|  | D3 | D6 | D9 | D3 | D6 | D9 |
| Chemical method | +109.77 | +108.6 | +112.7 | 5.23 | 5.16 | 5.08 |
| Biological method | +113.8 | +113.8 | +114.5 | 5.8 | 5.25 | 5.24 |

**Figure 1.**
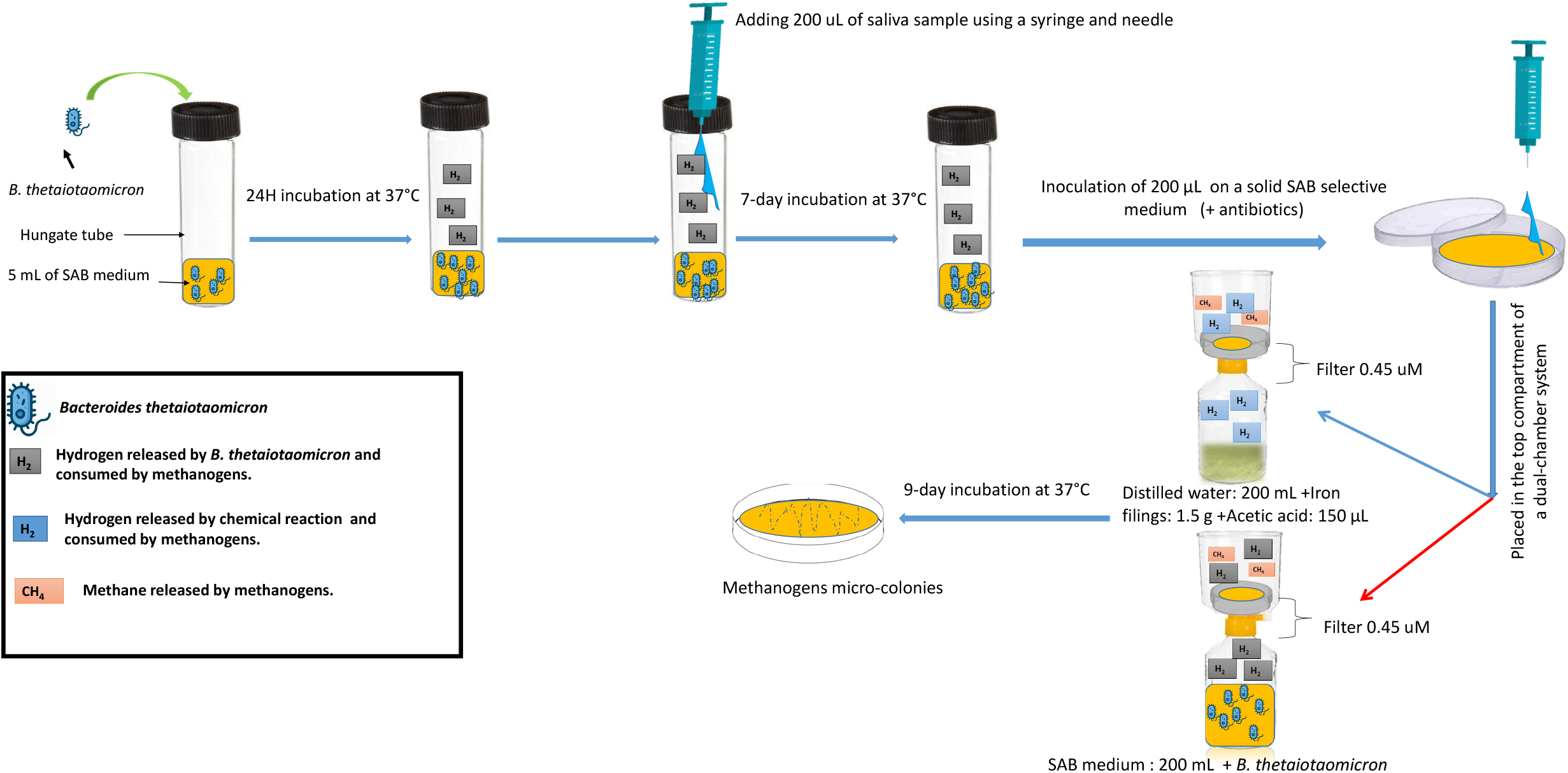
Aerobic culture of methanogens in a double-chamber system on agar plates using either a microbiological method for hydrogen production (red arrow) or a chemical method for hydrogen production investigated here (blue arrow).

In a second step, by culturing the reference methanogen strains in the presence of the negative and positive controls, we obtained hydrogen production in all tubes and observed a correlation between hydrogen production and the amount of iron filings used and the acid acetic concentration. The hydrogen production using the chemical method was maintained until day 5, and the average pH of the mixture was 4.09 ± 0.15 and 4.35 ± 0.23 in the SAB culture broth and distilled water, respectively. *M. oralis* colonies that were confirmed by PCR sequencing were visible on day 3 postinoculation using the chemical method (Supplementary Fig. 1). The negative control consisting of PBS inoculation instead of *M. oralis* remained negative.

To compare the chemical method with the biological method, *M. smithii* was inoculated in SAB agar plates using both methods. Using the chemical method, inoculation of SAB agar plates with *M. smithii* yielded colonies as early as day 3 postinoculation, whereas they appeared on day 9 using the biological method (Supplementary Fig. 1, Fig. 2). All negative controls remained negative. The colonies were confirmed using PCR sequencing and real-time PCR. The average number of colonies of *M. smithii* was significantly higher using the chemical method than the biological method, regardless of the day of follow-up (Student’s test, p <0.0001, p <0.001 and p = 0.0001 for day 3, day 6 and day 9 postinoculation, respectively) (Table 4). Moreover, cultures of these two strains were obtained in three days rather than in nine days using the biological method (17). In addition, the chemical method was easier to set up than the biological method because it incorporated distilled water instead of the SAB culture broth used in the chemical method, in contrast to the biological method in which only the SAB culture broth can be used (Supplementary Table 1). Additionally, the biological method requires maintaining a *B. thetaiotaomicron* culture in the exponential growth phase to release efficient hydrogen production, which is not the case for the chemical method.

**Figure 2.**
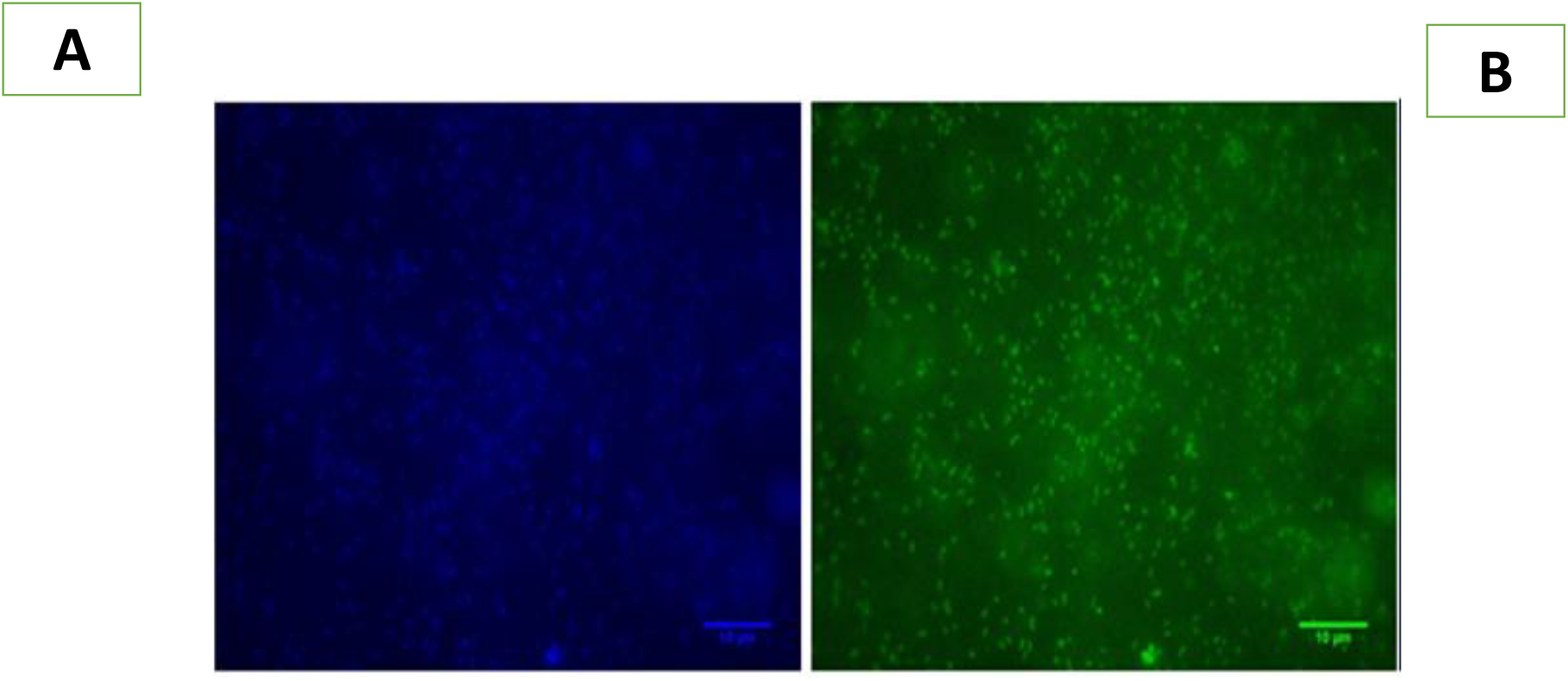
(A) Fluorescence emitted by *M. smithii* by applying the blue filter set. (B) Fluorescence emitted by *M. smithii* by applying the green filter set.

Finally, we compared the chemical method with the biological method for isolating methanogens from 50 saliva samples collected from 50 individuals after 9 days of incubation. The number of positive samples was 6/50 (12%) at day 3 postinoculation using both methods, and one additional saliva sample was positive at day 6, so that a total of 7/50 (14%) were positive using both methods. No additional sample was positive at day 9 postinoculation, regardless of the culture method. In these 7 culture-positive saliva samples, the average number of colonies at day 6 and day 9 postinoculation was significantly higher using the biological method than using the chemical method (Student’s test, p = 0.005 and p = 0.04), whereas there was no significant difference in the average number of colonies on day 3 (Student’s test, p = 0.09) (Table 4). Cultures from 6/25 (24%) of the samples from tobacco smokers yielded colonies as early as day 3 in both methods versus 1/25 (4%) of the samples from nonsmokers at day 6 in both methods. Colonies yielded autofluorescence in both methods (Fig. 2 and Fig. 3), whereas no colony or autofluorescence was observed in the negative controls. PCR sequencing and real-time PCR identified three *M*. oralis-positive and three *M. smithii-positive* samples in the tobacco smoker samples and one *M*. oralis-positive sample in one nonsmoker sample. Applying the chemical method for the isolation of *M. oralis* from saliva samples yielded significantly more colonies more rapidly than the biological method used in parallel. We hypothesized that the chemical method allowed mastering the kinetics of hydrogen production in such a way to optimize the atmosphere in the double-chamber system used in our experiments. These results show that the chemical method is more effective than the biological method for the rapid culture of methanogens.

**Figure 3.**
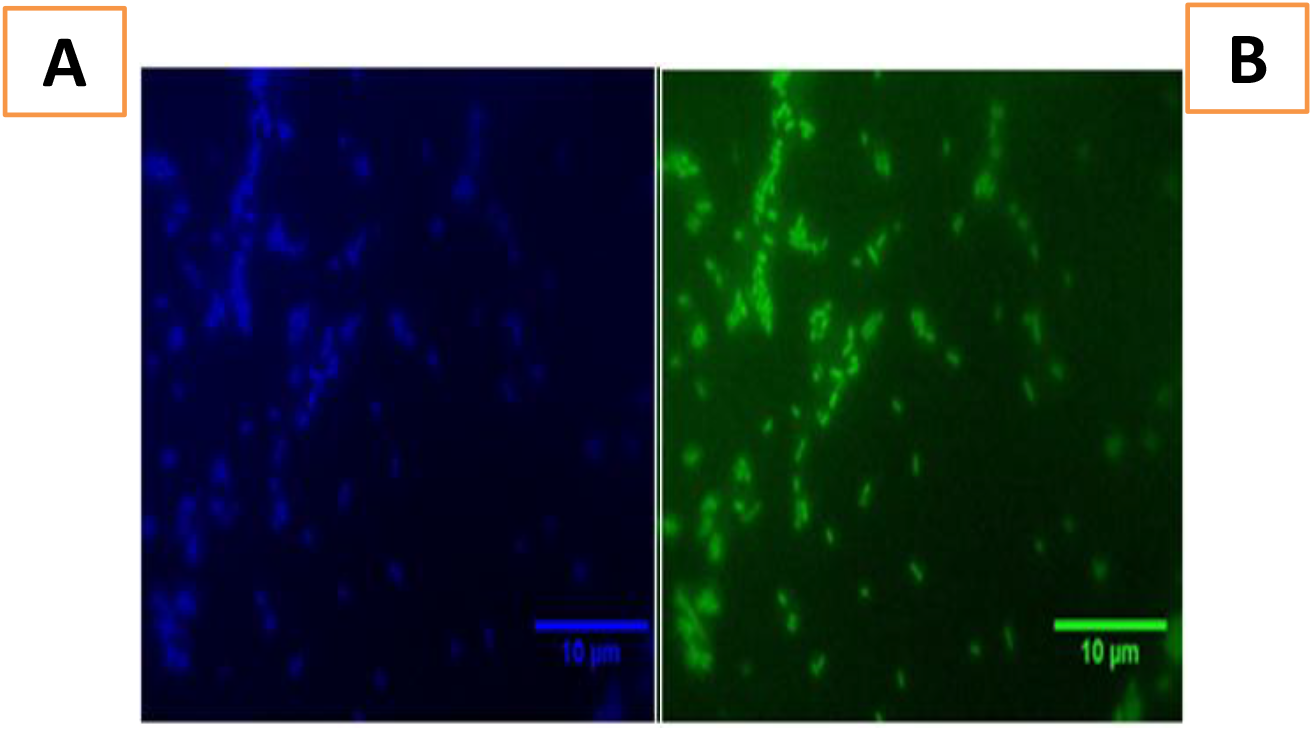
(A) Fluorescence emitted by *M. oralis* by applying the blue filter set. (B) Fluorescence emitted by *M. oralis* by applying the green filter set.

**Table 3.** Redox potential of the upper compartment.

Redox potential (mv) Chemical method Biological method
|  | D3 | D6 | D9 | D3 | D6 | D9 |
| --- | --- | --- | --- | --- | --- | --- |
| Distilled water | -28.1 | -81.2 | -108.9 | -4.5 | -66 | -91.6 |
| SAB medium | +13.5 | -116.5 | -77.5 | +19.1 | -7.1 | -34.9 |

**Table 4.** Comparison of the two methods according to the average of the colonies counted by ImageJ.

| Methods | Chemical method |  |  | Biological method |  |  | P-value |  |  |
| --- | --- | --- | --- | --- | --- | --- | --- | --- | --- |
|  | D3 | D6 | D9 | D3 | D6 | D9 | D3 | D6 | D9 |
| *M. smithii* | 377±72 | 659±168 | 1116±193 | 0 | 0 | 436±74 | <0.0001 | <0.001 | 0.0001 |
| Saliva samples | 190±90 | 509±133 | 700 ± 232 | 305±139 | 722±96 | 936±136 | 0.09 | 0.005 | 0.04 |
| Negative controls | 0 | 0 | 0 | 0 | 0 | 0 | NA | NA | NA |

NA : Not adapted

Accordingly, the observation that almost a quarter of the saliva samples grew *M. oralis* and *M. smithii* is in agreement with a previously published report (22). Interestingly, we confirmed that methanogens are more prevalent in the saliva from tobacco smokers than in the saliva collected from nonsmokers. These results are consistent with those found previously (22), in which there was also a predominance of *M. oralis* over *M. smithii* in saliva samples. These observations also confirm that *M. oralis* is the most prevalent methanogen in the oral cavity. We measured the redox potential in the culture using the chemical method to monitor whether this chemical reaction created anaerobiosis. Our results indicate that this chemical reaction is reducing and confirm that acetic acid is an efficient oxidizer of iron filings (20). The pH of the agar plates remained neutral throughout the inoculation period both in the chemical method and the biological method; this observation suggests that the acidity produced in the lower compartment of the double-chamber device in the chemical method does not reach the upper compartment.

The present study highlighted some advantages of the chemical method over the biological method (Supplementary Table 1). The chemical method is simple to set up, the hydrogen production is controlled so that the time of appearance of the first colonies is fast due to the immediate hydrogen production, and this chemical method can be done with both distilled water and SAB culture broth. Given the simplicity, speed and efficiency of this chemical method of hydrogen production, it could replace the biological method incorporating *B. thetaiotaomicron* in the isolation and culture of methanogens. Using the chemical method will ease the routine isolation and culture of methanogens in microbiology laboratories to ease and speed the isolation and culture of these opportunistic pathogens (1).

## ACKNOWLEDGEMENTS

COG and GG benefit from Ph.D. grants from the Fondation Méditerranée Infection, Marseille, France. This work was supported by the French Government under the «Investissements d’avenir» (Investments for the Future) program managed by the Agence Nationale de la Recherche (ANR, fr: National Agency for Research), (reference: Méditerranée Infection 10-IAHU-03). This work was supported by Région Sud (Provence Alpes Côte d’Azur) and European funding FEDER PRIMMI.

