## Supplementary figures and images for "Culture of salivary methanogens assisted by chemically produced hydrogen"

### Supplementary Figure 1

Supplementary figure 1

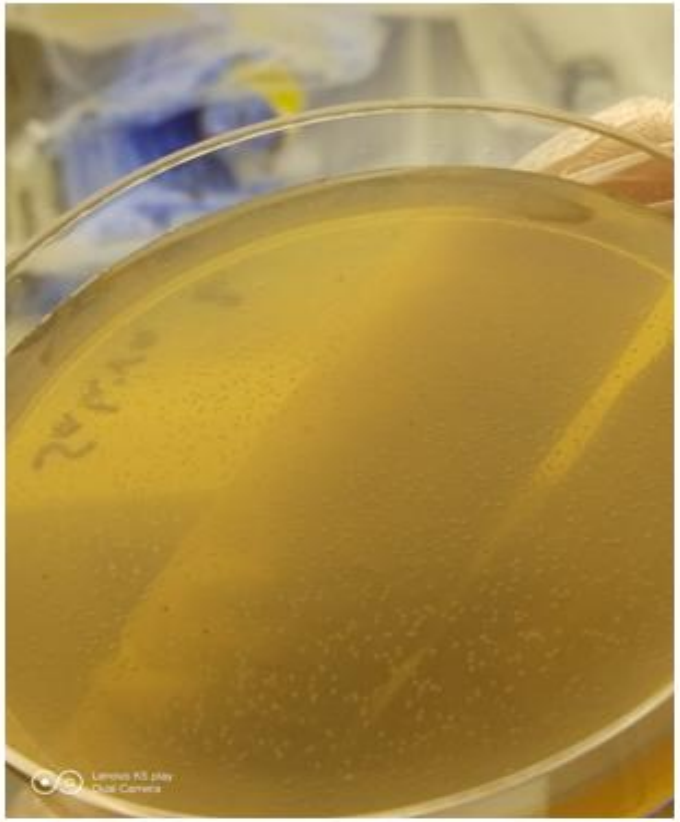

*Methanobrevibacter oralis*

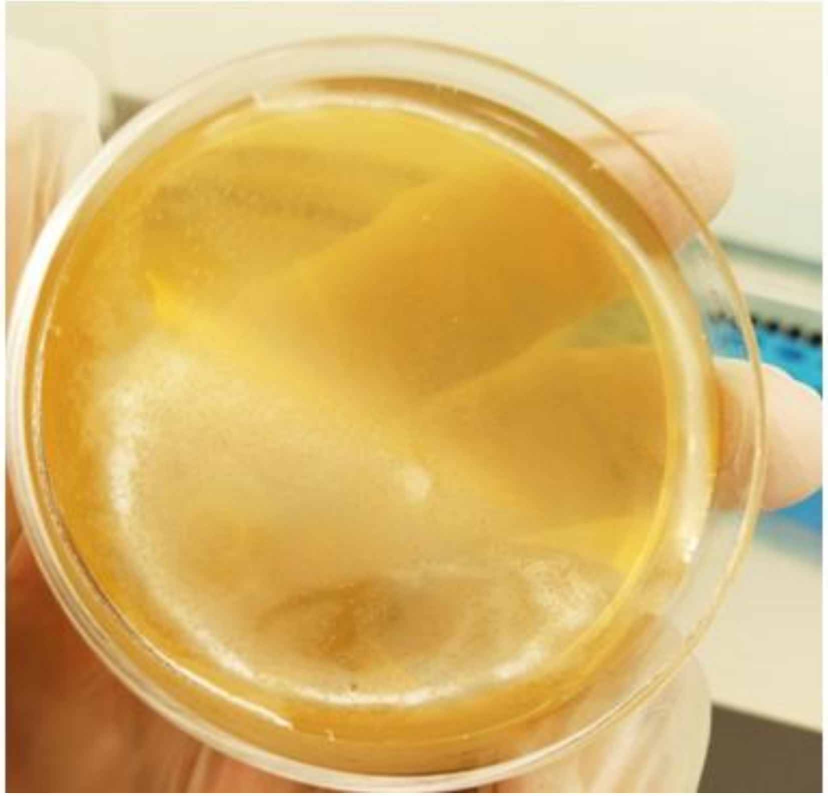

*Methanobrevibacter smithii*
